# Nextstrain automates real-time phylogenetic analysis of open data for endemic and emerging pathogens

**DOI:** 10.64898/2026.03.23.713807

**Authors:** Kimberly R. Andrews, Jennifer Chang, Cornelius Roemer, James Hadfield, Victor Lin, Anderson F. Brito, Richard Olumide Daodu, Isabel A. Joia, Kathryn Kistler, Allison Li, Louise H. Moncla, Miguel I. Paredes, Denise Kühnert, Laura Marcela Torres, Laura Voitl, Ivan Aksamentov, Emma B. Hodcroft, John Huddleston, John T. McCrone, John S. J. Anderson, Thomas R. Sibley, Jover Lee, Richard A. Neher, Trevor Bedford

**Affiliations:** Vaccine and Infectious Disease Division, Fred Hutchinson Cancer Center, Seattle, WA, USA; Biozentrum, University of Basel, Basel, Switzerland; Swiss Institute of Bioinformatics, Lausanne, Switzerland; Howard Hughes Medical Institute, Seattle, WA, USA; Instituto Todos pela Saúde, São Paulo, SP, Brazil; Center for Artificial Intelligence in Public Health Research, Robert Koch Institute, Berlin, Germany; Department of Mathematics and Computer Science, Freie Universität Berlin, Berlin, Germany; Department of Biology, University of Washington, Seattle, WA, 98195 USA; Department of Pathobiology, School of Veterinary Medicine, University of Pennsylvania, Philadelphia, PA, USA; Washington Department of Health, Shoreline, WA, USA; Swiss Tropical and Public Health Institute, University of Basel, Allschwil, Switzerland

## Abstract

**Motivation:** Genome sequencing provides an exceptional window into the evolutionary and epidemiological dynamics of endemic and emerging pathogens, and thus allows for better, more targeted, public health interventions. Online genomic surveillance platforms can provide near real-time insight into these dynamics.

**Results:** Nextstrain provides continually updated real-time genomic surveillance for 21 viruses and the bacterial pathogen *Mycobacterium tuberculosis*, with most analyses relying solely on open sequence data. Each pathogen includes steps to fetch and curate open data, classify sequences using established nomenclature systems, perform phylogenetic analyses, and share the results publicly. These analyses are automated, with most running daily to provide continually updated snapshots of pathogen evolution.

**Availability and Implementation:** All source code is available at https://github.com/nextstrain. Phylogenetic results can be visualized and downloaded at https://nextstrain.org/pathogens, and open sequence data and curated metadata are available at https://nextstrain.org/pathogens/files.

## 1 Introduction

The importance of rapid, open sharing of pathogen genome sequence data has become increasingly apparent over the last decade, with major outbreaks of multiple pathogens demonstrating the value of timely access to genomic data. Open-source genomic surveillance platforms, such as Nextstrain [1], CoV-Spectrum [2] and outbreak.info [3], leverage these data to perform continually-updated analyses and make the results immediately available to the public. This process rapidly transforms shared genomic data into insights regarding pathogen dynamics, including geographic spread and emergence of new variants. This understanding enables public health agencies, epidemiologists, academics, and the broader public to respond quickly and effectively to emerging public health threats.

Nextstrain provides continually-updated and highly customizable *real-time phylogenetic* analyses for multiple pathogens of public health significance. These analyses integrate epidemiological data into phylogenetic inference, where sampling dates are used to estimate time-resolved genealogies under a molecular clock and sampling locations are used to estimate phylogeographic history. This places our core analyses just past traditional phylogenetics in the realm of bare-bones phylodynamic inference [4]. We describe these analyses as “real-time” because they are kept fully up-to-date with available data, although analyses may be less up-to-date with respect to ongoing evolution, depending on the frequency at which new data becomes available. At the time of our initial publications describing these analyses, we supported workflows for influenza [5], dengue, Zika, and Ebola viruses [1]. Over time, Nextstrain has expanded these analyses to include a total of 21 viral pathogens and the bacterial pathogen *Mycobacterium tuberculosis*. Since 2020, these real-time analyses have provided critical biological and epidemiological insights that have informed public health responses to emerging pathogens SARS-CoV-2, mpox, Oropouche, and avian influenza, while also providing ongoing analysis of endemic pathogens such as seasonal influenza and RSV.

The expansion and maintenance of Nextstrain’s real-time analyses required the development of significant computational infrastructure. Here we describe the bioinformatics strategy that enables these analyses across multiple pathogens. Nextstrain’s pipelines are primarily based on *open data*, defined here as data that have been shared in a fashion that permits resharing and analyses with attribution to the original data generators. The exceptions are a subset of pipelines for SARS-CoV-2 and influenza, which are based on data from GISAID [6] and are thus restricted in re-sharing of curated data and analysis results. The open data pipelines include four main steps: 1) curation of open data using databases such as GenBank, Sequence Read Archive (SRA), and Pathoplexus; 2) quality control and clade calling; 3) customizable phylogenetic analyses; and 4) interactive visualization. These steps are automated, often running daily, and the outputs are made publicly available through nextstrain.org and via API access. The phylogenetic analysis step is highly flexible and can be tailored to run multiple different analyses for the pathogen of interest, including analyses focused on certain lineages, parts of the genome, geographic regions, or time periods. These analyses have direct surveillance utility, and can also serve as starting points for downstream analyses by public health agencies or academic labs.

Compared with our initial Nextstrain real-time analysis workflows [1], our current workflows retain the same core steps but are organized quite differently (see **Supp Text** for details). At the time of the 2018 publication, Augur was still a monolithic repository with per-pathogen configuration. Since then we have moved to a modular approach with Augur as a toolkit [7], with each pathogen analysis instantiated as its own workflow that strings together calls to Augur alongside bespoke scripts and other software. Beyond Augur, since 2018 we have improved the visualization capabilities of Auspice and added Narratives [8], introduced sequence quality control and clade-calling via Nextclade [9], added support for sharing analyses through GitHub or Nextstrain Groups, and added support for fully automated updates.

## 2 Materials and Methods

### 2.1 Overview of pipeline architecture

Nextstrain real-time genomic monitoring pipelines include steps to 1) ingest and curate sequence data and associated metadata from public repositories; 2) classify sequences to clades, lineages, or genotypes; 3) perform evolutionary analyses; and 4) share and visualize results at nextstrain.org (**Fig. 1**). These steps rely heavily on Nextstrain software packages, including Nextclade for viral sequence classification [9], Augur for phylogenetic analyses [7], Auspice for interactive visualization, and a RESTful API for data sharing. The pipelines are automated to run at regular intervals (usually daily) using GitHub Actions to keep the analyses up-to-date as new sequences become available. Sequence data and metadata are primarily fetched from GenBank, SRA, and Pathoplexus, since these are large, centralized, open databases that are accessible through APIs. However, the pipelines can be tailored to fetch sequence data from any source that provides API access. All of our pipelines are built using Snakemake workflow manager [10], and the code for each pipeline is publicly available on GitHub, with each pathogen having a separate repository under the broader Nextstrain organization. Within this general framework, our pipelines have a number of features that differ between viral and bacterial analyses, which we describe in the following sections.

**Figure 1.**
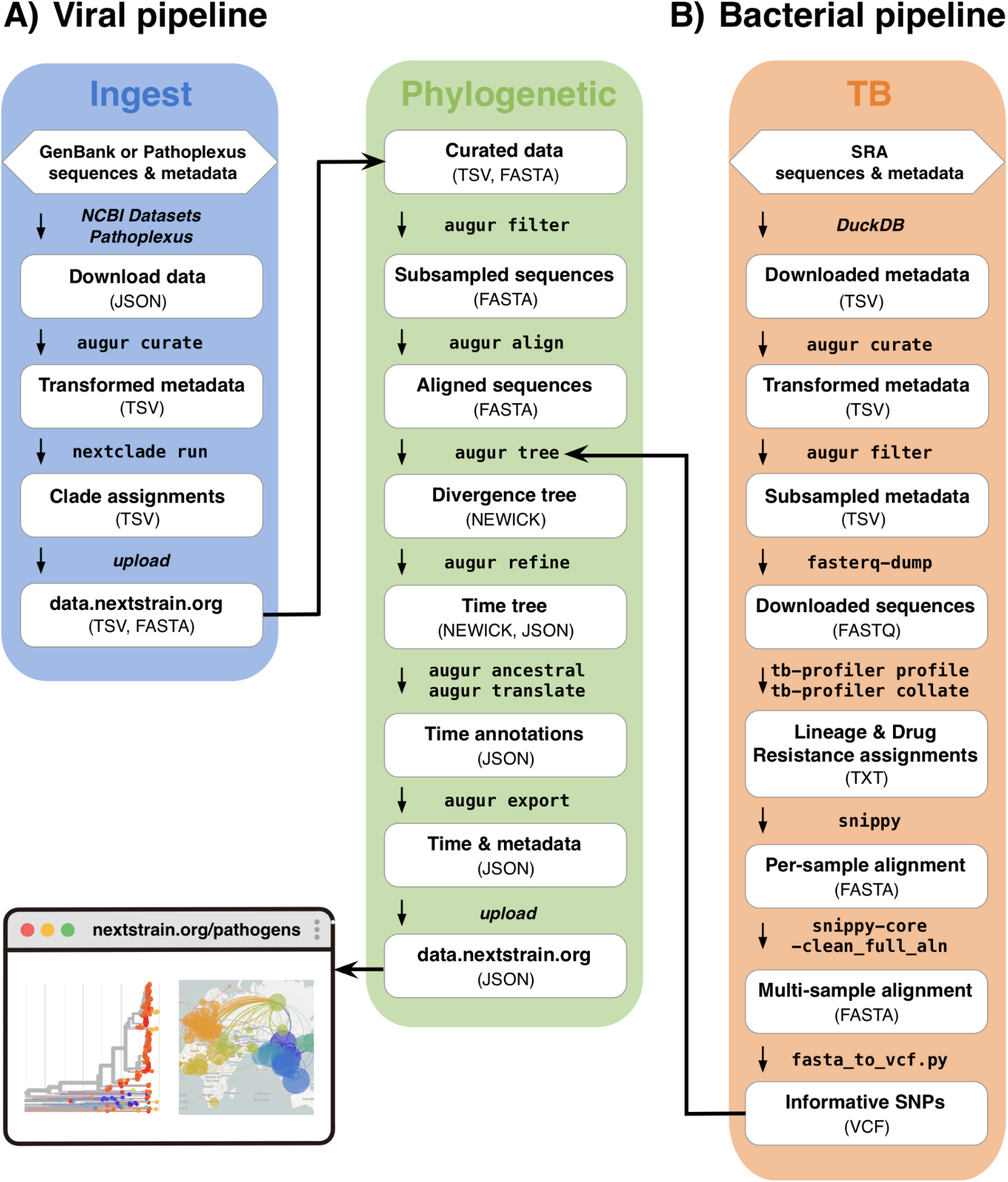
Outline of major steps for Nextstrain real-time analysis pipelines for viral and bacterial pathogens. Each pipeline is customized to the pathogen of interest, and the steps illustrated here represent those commonly shared across pipelines. The steps are arranged by the Snakemake workflows in which they occur within the pipelines, with each workflow shown as a colored block. The left hand arrow indicates that the outputs of the ingest workflow are used as inputs for the phylogenetic workflow of the viral pipeline. The right hand arrow indicates that the final steps of the bacterial pipeline are the same as those of the phylogenetic workflow of the viral pipeline; the arrow marks the point at which the two pipelines converge. The bacterial workflow is customized for *Mycobacterium tuberculosis*.

### 2.2 Viral analysis pipeline

For viral pathogens, our real-time analysis pipelines are divided into two Snakemake workflows called “ingest” and “phylogenetic” (**Fig. 1**). The *ingest workflow* fetches sequence data and metadata from external repositories, and then curates the metadata. In addition, if a Nextclade dataset is available for the pathogen of interest, the ingest workflow uses Nextclade to perform sequence quality assessment and assign lineages to each sample. The outputs of the ingest workflow are then used as inputs to the *phylogenetic workflow*, which performs subsampling, alignment, and a wide range of evolutionary analyses. In addition, our viral analyses include automation that runs the ingest and phylogenetic workflows on a regular basis to provide continually updated genomic monitoring. For some viral pathogens, our analysis pipelines also include a Snakemake workflow called “nextclade.” This workflow is not run as part of the continually updated real-time analyses, but instead generates a stable reference phylogeny that can be used as a Nextclade dataset. The Nextclade workflow is only updated after new lineages are designated or substantial new diversity has emerged [9] for the pathogen of interest. In the following sections we describe each of these analysis components in greater detail.

#### 2.2.1 Ingest workflow

The first step of the ingest workflow fetches all consensus genome sequences and associated metadata for the pathogen of interest (measles, SARS-CoV-2, etc…) from an external database. Most of our viral workflows currently fetch data from GenBank using the programs NCBI Datasets [11] or Entrez [12]. However, a growing number of our workflows fetch data from Pathoplexus [13]. The second step of the ingest workflow transforms the metadata to a format that is more tractable for downstream analyses and standardizes metadata values. For example, we transform collection dates to the ISO 8601 international standard format (YYYY-MM-DD), although with incomplete dates masked with “XX”, and parse geographic location information into standardized values that are separated into fields for country, division, and location. For viruses that have a Nextclade dataset, our ingest workflow also uses Nextclade to assign lineages and perform sequence quality assessment for each sample, and appends a column to the metadata file with the lineage assignments and quality control metrics for each sample. The final step of the ingest workflow uploads the sequences and transformed metadata to data.nextstrain.org to enable downstream phylogenetic analyses by Nextstrain workflows or external users (**Fig. 1**) The full list of sequence and metadata files that are publicly available for each pathogen can be viewed at nextstrain.org/pathogens/files.

#### 2.2.2 Phylogenetic workflow

The first step of the phylogenetic workflow downloads the outputs of the ingest workflow, which include the consensus genome sequences and the harmonized metadata. The subsequent steps are highly customizable to perform analyses addressing a wide range of research questions, and most of our pipelines perform multiple analyses per pathogen. These steps primarily use Nextstrain’s Augur software [7], which is a command line toolkit for pathogen phylogenetic analysis that provides built-in functionality for many data processing tasks, while also serving as a wrapper to run and standardize inputs and outputs for widely used bioinformatic tools such as MAFFT [14], IQ-TREE [15], and TreeTime [16]. Typically the first of these steps interrogates the metadata to select an appropriate subset of sequences to address the research question of interest. This subsampling step is highly customizable and includes options such as selecting or excluding sequences that match specified metadata parameters, selecting certain numbers or weighted proportions of samples across groups of metadata parameters, and setting a maximum number of total sequences. For example, a common approach is to subsample evenly over time and geography to obtain a representative worldwide subset of approximately 3000–5000 samples. We generally target a few thousand samples as this size of dataset gives sufficient data for understanding dynamics while staying performant in workflow construction and in Auspice JavaScript visualization. A few thousand viral sequences will run in roughly 1 hour on a laptop and the resulting Auspice visualization will load in less than 0.5 seconds (**Supp. Table 1**). Scaling to more than 10k or 20k tips will start to bottleneck at the tree building stage in terms of both CPU time and memory.

**Table 1.**
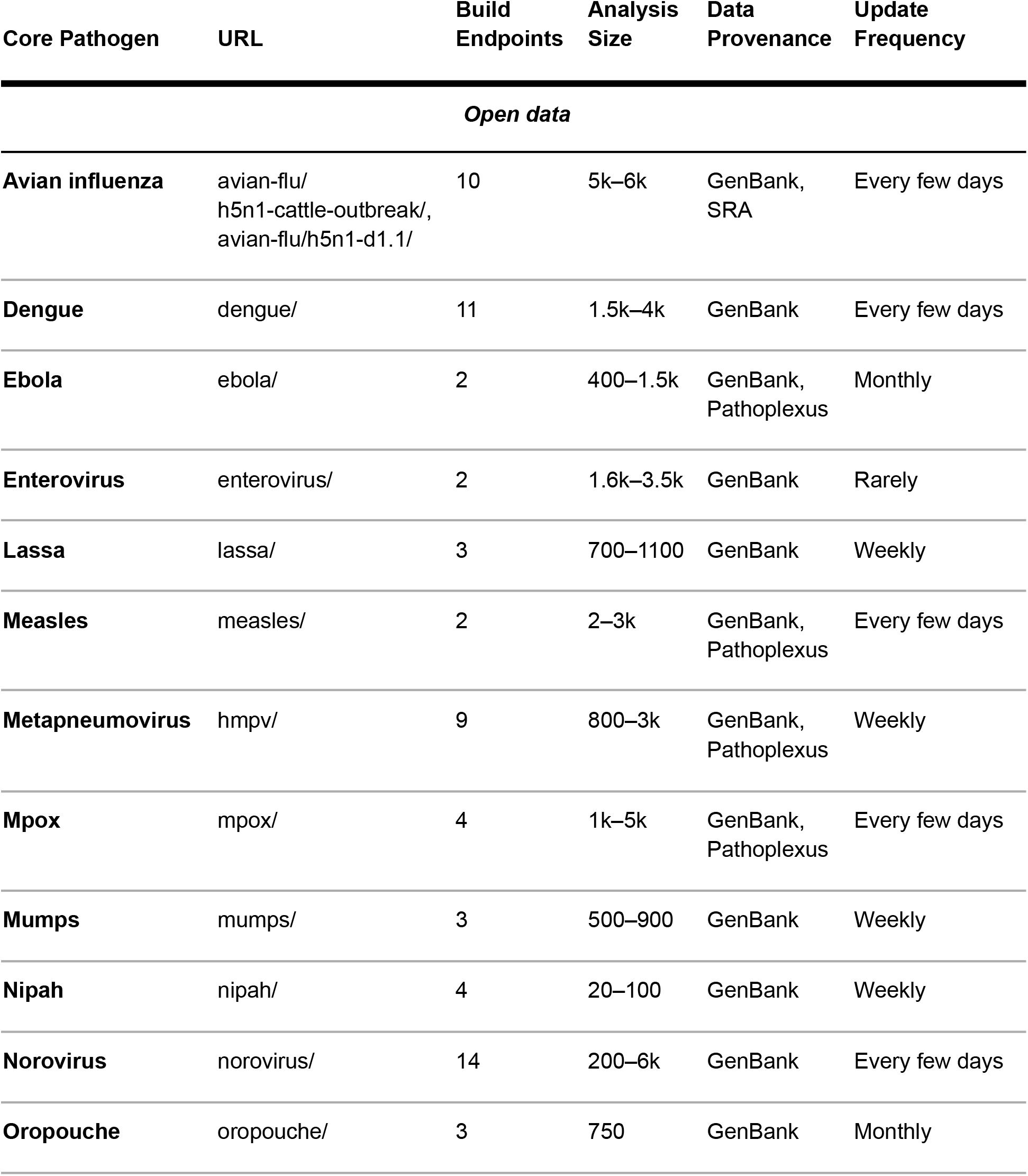
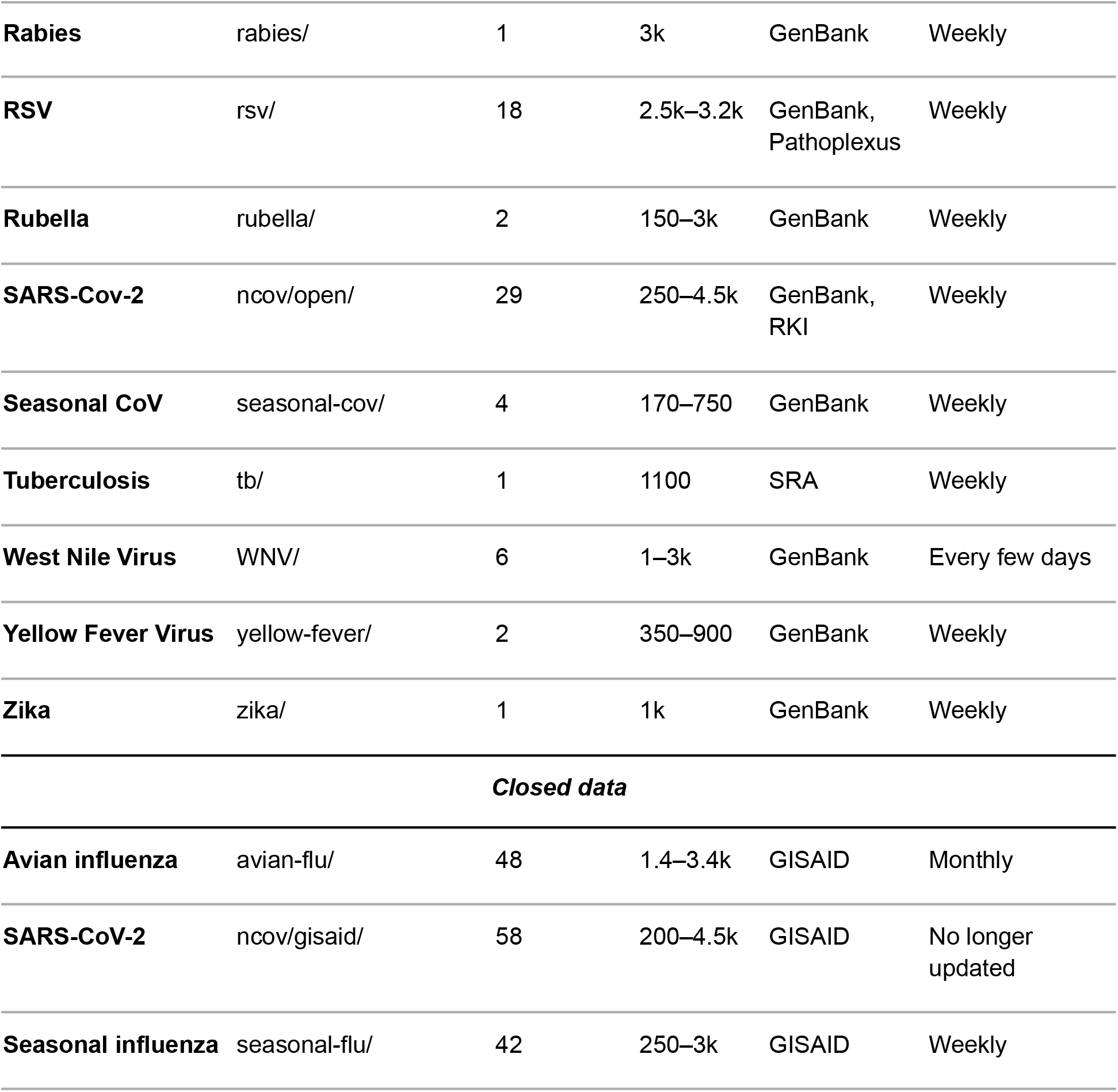
Summary of real-time analysis features for core pathogens. Build endpoints = number of phylogenies produced by the pipeline; analysis size = number of samples included in the phylogenies, shown as a range when multiple phylogenies are generated; update frequency = the approximate frequency at which the automated pipeline completes the phylogenetic workflow, which depends on how often the automated pipeline is initiated and how frequently new samples are deposited in repositories. Data for this table were updated on March 5, 2026.

After subsampling, the consensus genome sequences of the selected samples are then aligned to a reference genome, which is a representative genome sequence for the pathogen of interest; our primary criteria for selecting a reference genome are that it should be widely used and have high quality sequence data and annotations. The aligned sequences are then used to produce a maximum likelihood phylogeny with IQ-TREE and a time-resolved phylogeny with TreeTime. IQ-TREE uses a standard GTR model with command `-m GTR --ninit 2 -n 2` `--epsilon 0.05`. This is tuned to be fast and robust as branch lengths will be polished anyway by TreeTime in the following step. This call to IQ-TREE is configurable by users.

One of the main outputs of the phylogenetic workflow is a JSON file which contains information regarding the structure of the phylogenies, node annotations such as nucleotide and amino acid mutations, and relevant metadata for each sample. This file can be used as input to Nextstrain’s Auspice software for interactive visualization of the phylogenies. The final step of the phylogenetic workflow uploads this JSON file to nextstrain.org, where it can be publicly viewed with Auspice.

#### 2.2.3 Automation

We automate ingest and phylogenetic workflows to run at a particular cadence, generally once daily. This automation is effected through a GitHub Actions workflow that performs the following steps: 1) run the ingest workflow to download and curate data from an external repository; 2) check whether any new sequence data has been deposited in the external repository since the last run of the ingest workflow by comparing file identifiers between the previous and new download; 3) run the phylogenetic workflow only if new sequence data has been detected; and 4) post the results to nextstrain.org. These steps ensure that the computationally expensive phylogenetic analyses only run if new sequence data are available.

For most pathogens, this GitHub Action runs daily, and analyses remain within the standard computational resource limits provided by GitHub Actions. For pathogens with large volumes of available sequence data and requiring multiple phylogenetic analyses (SARS-CoV-2, influenza, mpox, RSV), we use GitHub Actions to launch workflows on AWS Batch for access to greater computational resources. To reduce computational load, these analyses are typically run weekly rather than daily. These automated processes produce continually updated resources including curated data and phylogenetic datasets available for online visualization, and for download as JSON files (**Fig. 2**).

**Figure 2.**
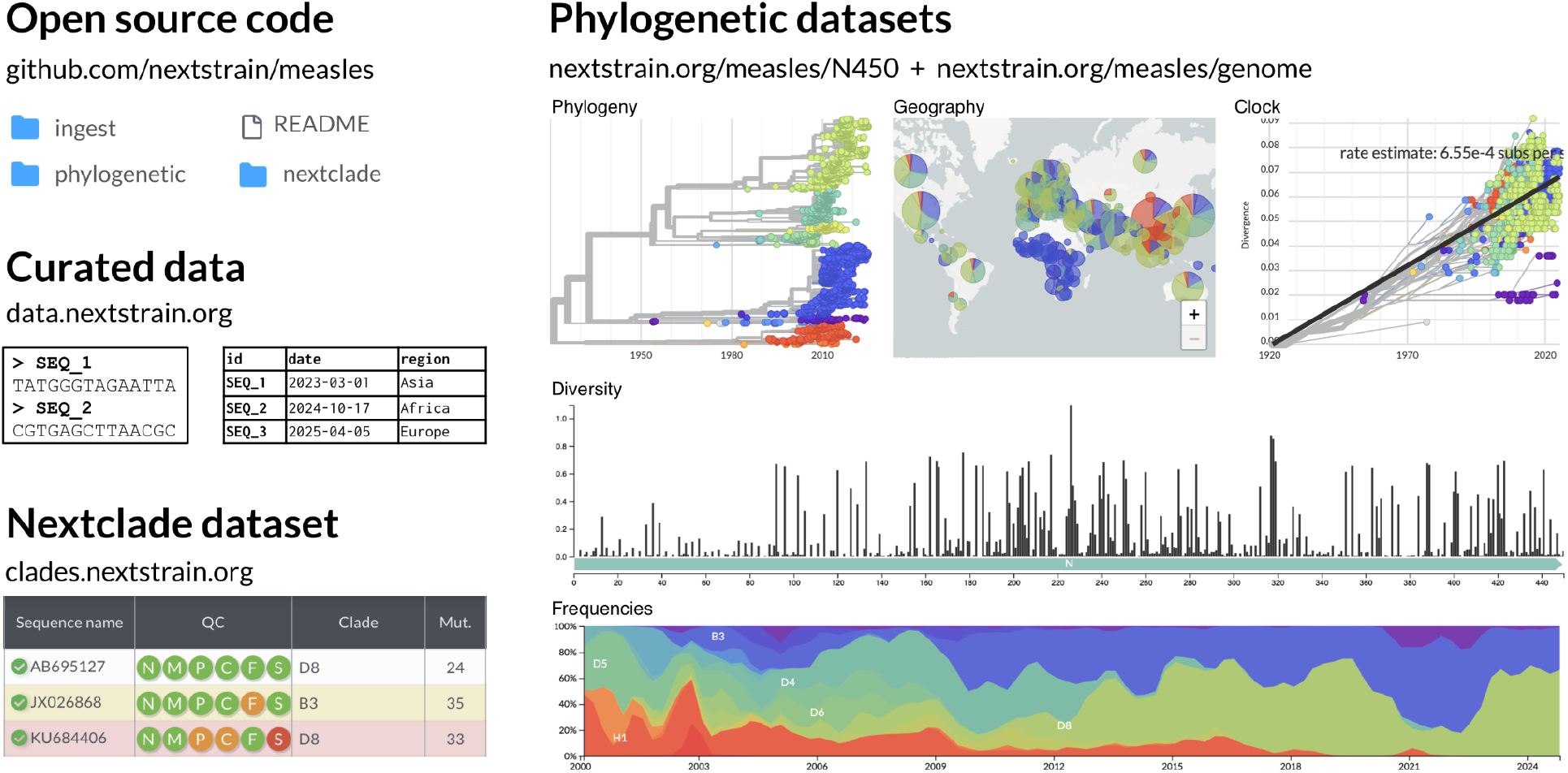
Example output resources for the real-time phylogenetic analysis of measles. These resources include 1) open source code for the analyses, available on GitHub; 2) sequence data and curated metadata from public repositories for the pathogen of interest; 3) a reference phylogeny that can be used as a Nextclade dataset; and 4) analytical results that can be visualized and downloaded at https://nextstrain.org.

#### 2.2.4 Nextclade workflow

Nextclade datasets are pathogen-specific tools that align viral genomes to a reference genome, perform sequence quality assessment and clade assignment, and identify mutations relative to the reference and clade founders[9]. An integral part of a Nextclade dataset is a reference phylogeny that includes representative samples from each clade, and the *nextclade workflow* generates these reference trees (**Fig. 2**). The nextclade workflow is similar to the phylogenetic workflow in our real-time analyses, but has a subsampling step that focuses on obtaining a subset of complete, high quality genomes that are representative of the genetic diversity for the pathogen of interest. Since the nextclade datasets serve as versioned and stable resources, this workflow is not run as part of the automated analyses, but can be triggered on demand.

#### 2.2.5 Customization for virus-specific features

Although our real-time viral pathogen pipelines follow the standardized format described in the preceding sections, each pipeline is customized to accommodate the biological features and research questions relevant to each virus. For viruses with more sequence data available for certain genes or loci than for whole genomes, we produce gene- or locus-specific phylogenies as well as whole genome phylogenies (e.g., E gene for dengue, N450 for measles). For segmented viruses (e.g., Lassa, Oropouche, influenza), we build separate phylogenies for each segment of the genome rather than whole genome phylogenies, since segment reassortment strongly impacts phylogenetic inference. In addition, many of our pipelines include analyses that use customized subsampling strategies to focus on certain geographic regions, lineages, or time periods. For example, when outbreaks occur, we often add a phylogeny focused on the lineage, geographic region, and time period associated with the outbreak (e.g., mpox, avian influenza). Also, many pipelines include customized coloring options to highlight important metadata or evolutionary outcomes (e.g., host species for West Nile Virus and rabies; epitope mutations for influenza).

### 2.3 Bacterial analysis pipeline

The pipeline for the bacterial pathogen, *M. tuberculosis*, shares the same basic steps and outputs as the viral pipelines. However, many of these steps are implemented differently to accommodate the biological differences between bacteria and viruses, such as the much larger genome sizes for bacteria (e.g., 4.4 MB for *M. tuberculosis*) (**Fig. 1**). The *M. tuberculosis* pipeline is also structured to include just one Snakemake workflow to run the entire analysis, rather than including separate ingest and phylogenetic workflows, as in the viral pipelines. In addition, the automation pipeline includes a number of differences to accommodate the greater computational resources needed to run the *M. tuberculosis* analysis. In the following sections, we describe the architecture of the *M. tuberculosis* pipeline, highlighting the differences from our viral pipelines.

#### 2.3.1 Ingest and phylogenetic workflow

One of the main differences of the *M. tuberculosis* pipeline compared to the viral pipelines is that it starts with raw short-read sequence data rather than consensus genome sequences. As a result, it fetches sequence data and metadata from the SRA database, whereas the viral pipelines fetch data from GenBank (**Fig. 1**). The pipeline starts by fetching the metadata for all *M. tuberculosis* samples with Illumina shotgun sequence data. In contrast to our viral pipelines, this step does not fetch any of the sequence data, due to the large file sizes for raw sequence data and large total number of *M. tuberculosis* samples on the SRA. Instead, the pipeline curates the metadata as described previously for the viral workflow, interrogates the transformed metadata to select a representative subset of approximately 1000 samples (rather than 3000-5000 samples typically used for the viral pipeline), and then fetches the corresponding read-level sequence data for only those samples (contained in FASTQ files). The smaller subset of samples is chosen for this pipeline due to the much larger numbers of mutations for *M. tuberculosis* compared to viral genomes, which can make visualization in Auspice slow for thousands of samples.

Next, the pipeline uses Snippy (https://github.com/tseemann/snippy) to align the sequence reads to a reference genome, identify variable sites for each sample, and create a multi-sample alignment of all the genome sequences. The pipeline also implements quality control checks at two different steps to remove samples with low-quality sequence data, and performs masking of sites in the alignment that are known to be difficult to genotype in *M. tuberculosis* [17] by replacing nucleotides at those sites with ambiguous bases (N’s). Next, to reduce the computational time required for downstream phylogenetic analysis, the information in the masked multi-sample alignment file is transformed into a compact VCF file that contains a summary of genotypes for all samples at phylogenetically informative sites. This file is used to build a maximum likelihood phylogeny using an Augur function that allows VCF files to be used as input to IQ-TREE.

Our *M. tuberculosis* pipeline also uses the program TBProfiler [18] to predict resistance to anti-tuberculosis drugs for each sample. This is accomplished by comparing the genome sequences of each sample against a database of mutations associated with drug resistance published by the World Health Organization and other sources [19]. TBProfiler also assigns a phylogenetic lineage to each sample using a reference database of lineage-specific mutations.

As in the viral workflow, the final output of the *M. tuberculosis* pipeline is a JSON file that can be used for interactive visualization with Auspice. The final step of the *M. tuberculosis* pipeline uploads this JSON file to nextstrain.org so that the results can be viewed publicly.

#### 2.3.2 Automation

The *M. tuberculosis* pipeline requires substantially more computational resources than most of our viral pipelines because it stores and analyzes Illumina shotgun sequencing FASTQ files, which are much larger than the FASTA files containing consensus genome sequences that are used in our viral pipelines. Thus, our automation pipeline for the *M. tuberculosis* analysis incorporates several features to accommodate this higher computational demand. First, the pipeline uses GitHub Actions to run the workflow with AWS Batch for larger instances with more CPUs, memory, and disk space. Second, every time we run the analysis, we cache the Snippy and TBProfiler results for each sample in an AWS S3 bucket, and then if future runs select any samples that were analyzed in a previous run, we download the Snippy and TBProfiler results for those samples from the S3 bucket rather than re-running those analyses. Since each run of the workflow selects approximately 1000 samples from all *M. tuberculosis* samples in the SRA that have Illumina shotgun sequence data (currently about 170,000 samples total), the cache builds up slowly over time, resulting in decreased workflow runtime as more runs are performed. Finally, we also address the higher computational demands of the *M. tuberculosis* pipeline by running the workflow only once a week, rather than once a day as for most of our viral analyses.

### 2.3.3 Customization for bacteria-specific features

Generalizing this pipeline to other bacterial species would require modifications to accommodate their distinct biology. Unlike the largely clonal *M. tuberculosis*, many bacteria undergo substantial homologous recombination, which violates the assumption of vertical inheritance underlying phylogenetic inference. Recombinant regions can be detected and masked with tools such as Gubbins [20]] or ClonalFrameML [21], though these methods assume recombination is a modest deviation from a clonal frame and are less suitable for highly recombinogenic species. Many bacteria also have open pangenomes with extensive gene gain and loss, so mapping to a single reference genome captures only part of the genome, and core- or pangenome-aware alignment strategies may be needed. Finally, the masking, lineage-assignment, and drug-resistance steps are all species-specific.

### 2.4 Visualization of results

The results for all of our real-time analyses are publicly viewable with Auspice at nextstrain.org. Each pathogen generally includes an interactive phylogenetic tree, a map of the world with pie charts showing the geographic distribution of samples, a plot showing the diversity of genetic variation at each position along the genome, and a plot showing the frequency of different types of samples over time. These plots are highly interactive; for example, users can zoom into certain parts of the tree, change the coloring in the plots to visualize different metadata parameters, view results for a single nucleotide position, view inferred ancestral states at internal nodes such as mutations or geographic region, filter sequences based on various metadata parameters, and many other options. Different components of the visualization are tightly integrated so that, for example, zooming into a part of the phylogeny will update the other panels to reflect this subset of the data, and changing a coloring option will change the coloring in all panels. The Auspice visualization also credits the individuals who contributed the open genomic sequence data by displaying their names in several ways. First, the names of individuals who submitted sequences to the public repository can be viewed by clicking on samples at the tips of the branches of the phylogeny, and samples can also be filtered based on submitter names. In addition, a text file with submitter names for each sequence included in the phylogeny can be downloaded at the bottom of the Auspice visualization page.

### 2.5 Modifying pipelines for custom analyses

In addition to providing real-time genomic surveillance, our pipelines also function as starting points for external users to develop new analyses that address their own research questions. Nextstrain provides multiple tools to streamline installation and usage of our pipelines, including a command line interface (CLI) and Docker, Conda, and Singularity (Apptainer) runtimes. In addition, our pipelines are highly customizable to address a wide variety of questions. For example, the pipelines can be modified to focus on certain phylogenetic lineages or localized geographic regions by modifying the subsampling parameters. Alternatively, users can modify the pipelines to use their own private sequence data and metadata, or a combination of private data and subsampled public data, and to enable coloring by any metadata parameter of interest. Furthermore, our pipelines can be modified for use with other pathogens (e.g., [22]). Most of our viral workflows can run on a laptop, and the use of GitHub Actions or AWS is not required.

However, workflows can be run on high-performance computing (HPC) or cloud environments to streamline analyses of large amounts of genomes (SARS-CoV-2) or more complex workflows (*M. tuberculosis*). Self-hosted runners can be connected to GitHub Actions to facilitate these use cases (internally, we use AWS Batch in these more demanding circumstances).

### 2.6 Sharing analysis results

Nextstrain also provides multiple methods for external users to share the results of their custom analyses publicly or privately. Users can visualize phylogenies by simply dragging and dropping JSON output files and metadata files onto the web-based tool https://auspice.us. Alternatively, Nextstrain provides two methods by which external users can visualize their results through nextstrain.org. First, any JSON output files that are stored on GitHub can automatically be viewed using nextstrain.org/community. Second, external users can set up a “Nextstrain Group” which is managed by Nextstrain and allows sharing of results at nextstrain.org/groups either publicly or privately within a group of researchers. Nextstrain also enables users to create interactive, data-driven narratives through the “Nextstrain Narrative” feature, which uses Auspice to display explanatory text alongside a corresponding view into the data that automatically updates as the user moves through the narrative [8].

## 3 Results

### 3.1 Summary of pipeline features

Nextstrain currently maintains analyses for 22 core pathogens listed in Table 1. Of these, 19 represent pathogens that have automated real-time analysis of open data, including 18 viruses and one bacteria. Two additional pipelines are not fully automated (Ebola and enterovirus D68) and one only uses restricted data (seasonal influenza). For two pathogens (avian influenza and SARS-CoV-2), a subset of analyses use restricted data (Table 1). The number of phylogenies produced by each of our pipelines ranges from one (for Zika, rabies, *M. tuberculosis*) to 87 (for SARS-CoV-2), with most pipelines producing between two and ten phylogenies (Table 1). Pipelines that generate multiple phylogenies typically do so to analyze multiple genomic regions (such as segments, genes, or loci) or to focus on multiple biologically important lineages, geographic regions, or time periods. All source code for the pipelines is available at https://github.com/nextstrain and analysis results are available at nextstrain.org/pathogens.

### 3.2 Examples of rapid outbreak responses

Over the last decade, Nextstrain real-time analyses have provided important insights into pathogen biology and epidemiology. The utility of these analyses has been particularly apparent during pathogen outbreaks, when rapid delivery of relevant information is critical for public health decision-making.

#### 3.2.1 Mpox

In July of 2022, a global outbreak of mpox virus resulted in the declaration of a public health emergency of international concern (PHEIC) [23], and researchers and public health workers from around the world began sharing mpox genome sequences on GenBank. In response, Nextstrain quickly developed a real-time analysis pipeline to conduct phylogenetic analyses using these sequences (**Fig. 3**). This pipeline required customization to accommodate several features of the mpox genome that differ from common epidemic viruses, including a much larger genome size (~200k base pairs) and the presence of large deletions and repetitive regions [24]. These features were addressed by masking repetitive genomic regions and adjusting parameter values for alignment of sequences to the reference genome. Results from this pipeline provided timely insight into mpox transmission dynamics [25] and facilitated a nomenclature system for mpox that includes an open method for proposing new lineage designations [26], [27]. In addition, this pipeline enabled Nextstrain to quickly respond to a second PHEIC declared for mpox in 2024 [28] by developing an additional phylogenetic analysis focused on a clade that was spreading rapidly in Central Africa. These mpox resources provided situational awareness and supported downstream genomic analysis by public health and academic groups [29–31].

**Figure 3.**
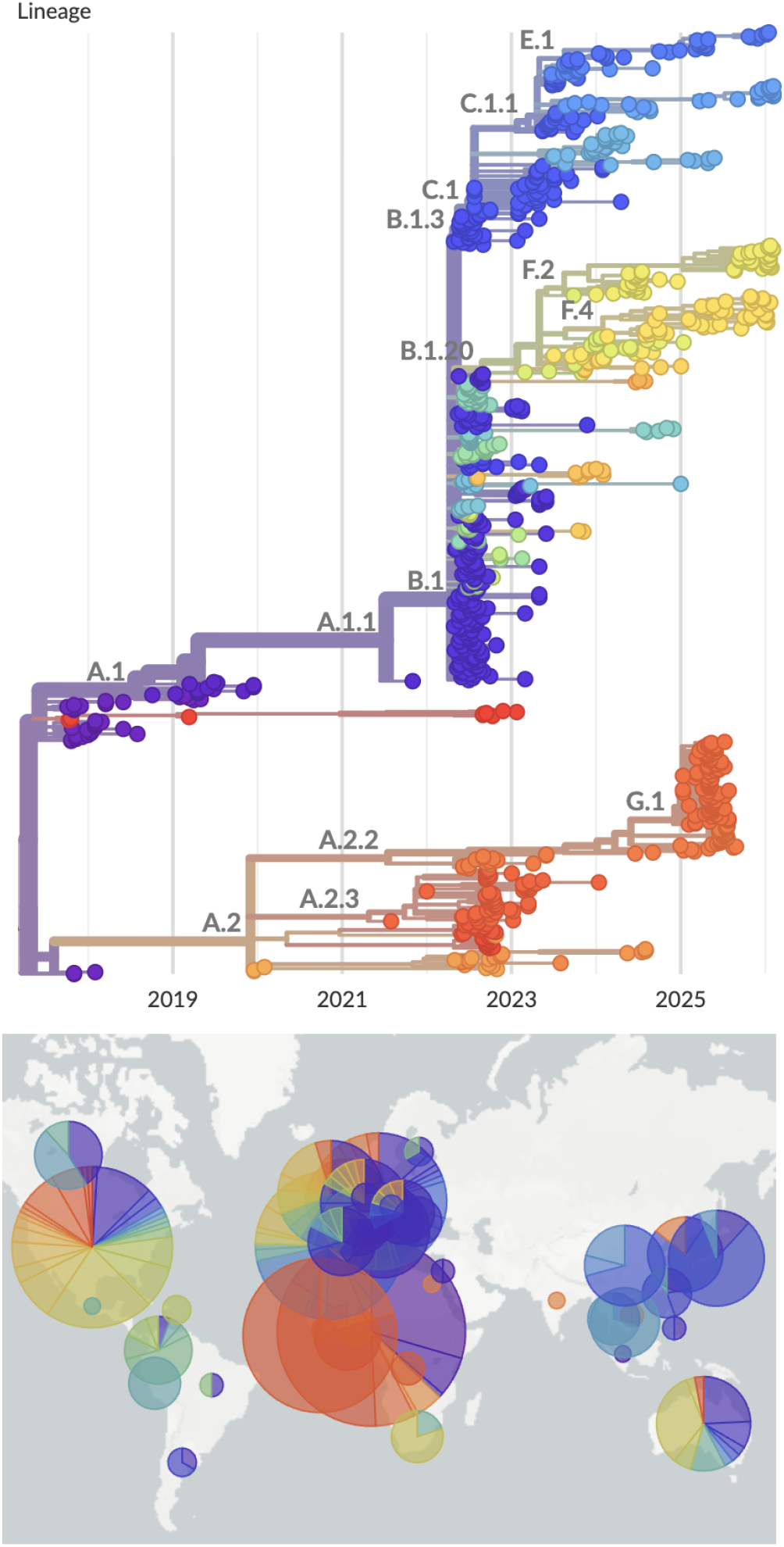
Time-resolved phylogeny of mpox clade IIb showing 927 viruses colored by lineage along with their geographic distribution. Branches shown have APOBEC3 mutation patterns distinct from ancestral branches [24], and so this is believed to represent human-to-human transmission starting in late 2017. A live version of this analysis is viewable at nextstrain.org/mpox/clade-IIb. Screenshot is from March 9, 2026.

#### 3.2.2 Avian influenza

When an outbreak of highly pathogenic avian influenza subtype H5N1 occurred in dairy cattle in North America in early 2024 [32], a Nextstrain real-time analysis pipeline was already in place for this pathogen, and we quickly expanded this pipeline to include new phylogenetic analyses focused on the cattle outbreak (**Fig. 4**). Sequence submission to public repositories was primarily by the National Veterinary Services Laboratories (NVSL) of the Animal and Plant Health Inspection Service (APHIS) of the U.S. Department of Agriculture (USDA). These submissions included both raw sequence data deposited to the SRA, and the corresponding genome assemblies deposited to GenBank, although genome assemblies were largely delayed in appearing on GenBank, with an average of 41 days between SRA and GenBank deposition. To address this issue, the Andersen Lab at Scripps Research developed an automated pipeline to assemble consensus genomes from SRA data and make the resulting genome assemblies publicly available through GitHub (https://github.com/andersen-lab/avian-influenza). In response, Nextstrain tailored its analysis pipeline to use consensus genome sequences from both GenBank and the GitHub repository. The phylogenetic analyses implemented in this pipeline provided rapid insight into the number and timing of spillover events between birds, cattle, humans, and other mammals; transmission dynamics between U.S. states; and potential evolutionary adaptations of the virus to mammalian hosts. Early in the outbreak, these analyses showed clear support for a single spillover from birds followed by transmission among cattle, highlighting a novel epidemiologic event that was distinct from outbreaks in poultry. As the outbreak in cattle grew and spread, these analyses allowed rapid identification of spillbacks of H5N1 from cattle into domestic poultry and cats, and the ability to discern the source of infections stemming from exposure to raw milk and raw pet food.

**Figure 4.**
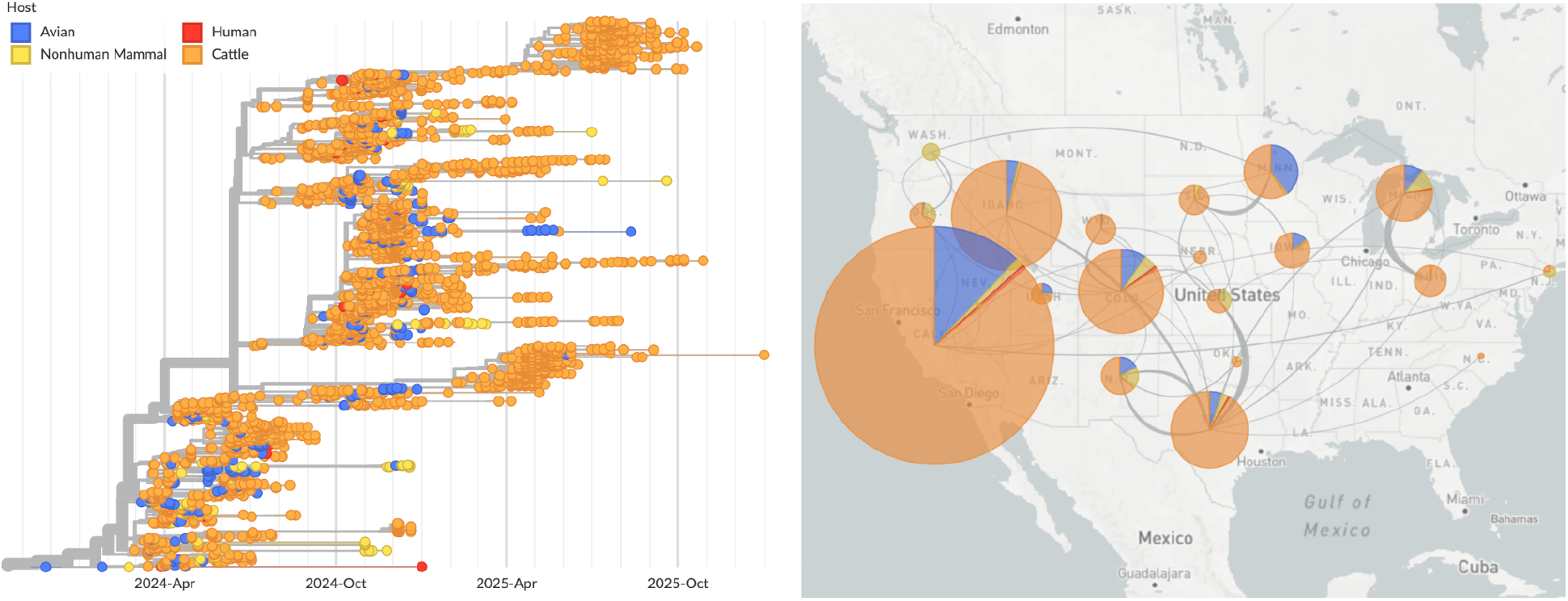
Full genome time-resolved phylogeny of avian influenza H5N1 outbreak in North America colored by host. A live version of this analysis is viewable at nextstrain.org/avian-flu/h5n1-cattle-outbreak/genome. Screenshot is from March 9, 2026.

### 3.3 Broader phylogenetic analyses

Beyond rapid outbreak response, analyses in Nextstrain provide an overview of the evolution and epidemiology of a number of endemic and epidemic pathogens. For example, these analyses have provided information regarding the number, timing, and geographic source of introductions of pathogens into different geographic regions. This information is important for developing effective prevention and control measures and is particularly relevant for epidemics and outbreaks of measles, mumps, Ebola, Zika and Oropouche (e.g., [33,34]). Nextstrain analyses have also been used to detect pathogen spillover to new host species, which provides important information regarding the probability of future spillover events and whether those events are likely to result in sustained transmission in the new host species (e.g., [35]). These spillover analyses are particularly relevant for rabies, avian influenza and Yellow Fever virus. Nextstrain has been used to analyze patterns of reassortment and recombination [36,37]. In addition, evolutionary analyses track the emergence and spread of new clades that have fitness advantages, and identify amino acid changes that may be responsible for those advantages [38]. These analyses can identify variants of potential biological relevance, and can improve decision making regarding strain selection for vaccine development in pathogens that undergo antigenic evolution [39].

## 4 Discussion

Nextstrain real-time analyses demonstrate the valuable insights that can be gained by pathogen genomic surveillance through the integration of open sequence data, open-source analytical software, and openly shared results. By combining these resources, Nextstrain now provides continually updated, publicly accessible genomic monitoring using open data for 19 important pathogens. These analyses provide up-to-date information regarding transmission dynamics, geographic spread, emergence of new variants, spillover to new host species, and other important evolutionary and epidemiological processes. This information is critical for detecting and responding to public health threats in a timely and effective manner, and these types of analyses have become indispensable parts of modern infectious disease research and outbreak response.

Nextstrain’s real-time analyses also illustrate the importance of making open pathogen genomic surveillance resources easily accessible. For example, genomic sequence data are most accessible from centralized databases that provide API access, allowing any user to retrieve data programmatically; such access is critical for automating Nextstrain’s real-time analysis pipelines. In contrast, databases that are password-protected or that require manual downloads through graphical interfaces need human intervention every time the analysis is updated, thereby hindering access, automation, and real-time analysis. For this reason, Nextstrain’s pipelines primarily rely on GenBank, SRA, and Pathoplexus databases, since these large, open, centralized repositories provide API access.

Effective genomic surveillance also relies on accessibility of software and analysis results. Nextstrain provides API access allowing any user to download the real-time analysis outputs, including open sequences, curated metadata, and phylogenetic analysis outputs. Nextstrain also seeks to maximize accessibility by making its computational infrastructure easy for external users to install, use, and customize. In addition, Nextstrain fosters collaboration by providing multiple methods for users to share their analysis results either publicly or privately. By increasing accessibility, these features empower local public health agencies and academic laboratories to develop customized analyses that can inform responses within their own local regions. For example, Nextstrain analyses have supported genomic surveillance of mpox in King County, Washington [30], SARS-CoV-2 in Pakistan [40] Ebola in the DRC [41] and Lassa in West Africa [36]. Several public health and research institutions further maintain their own continually updated analyses as Nextstrain Groups, including the Washington State Department of Health (nextstrain.org/groups/wadoh) and the Institut National de Recherche Biomédicale in the Democratic Republic of the Congo (nextstrain.org/groups/inrb-mpox).

Open-source genomic surveillance platforms rely on the generous sharing of sequence data, and in turn these platforms can increase the visibility and impact of that data. With Nextstrain, we aim to ensure that data generators benefit from sharing — by increasing the reach and utility of their work while giving clear credit to the individuals behind it. GenBank, SRA, and Pathoplexus all credit individuals who submit sequence data by including submitter names in associated metadata, though the prominence of this credit and the terms attached to data reuse vary among repositories. Pathoplexus differs from GenBank and SRA in that it couples open resharing with explicit terms of use and includes both open and restricted sequence data;restricted data may be used in unpublished work like Nextstrain real-time analyses, but not in scientific publications or preprints. To surface this credit as broadly as possible, the curated metadata generated by our ingest workflow includes submitter names for each sequence, along with restriction status and terms of use for Pathoplexus sequences. Submitter names and restriction status for each sequence can also be viewed and downloaded directly from the Auspice visualization at nextstrain.org.

Although we focus here on Nextstrain’s real-time analyses, Nextstrain is just one resource within a broader ecosystem of valuable open-source pathogen genome surveillance resources. Like Nextstrain, many of these resources are highly flexible, which often allows them to be used synergistically. For example, Nextstrain’s Auspice visualization tool provides functionality for exporting and viewing phylogenetic trees in other open-source software including Taxonium [42] and MicrobeTrace [43]. Conversely, the open-source program UShER [44], which rapidly places new genome sequences on very large viral phylogenies via matOptimize [45], provides options to view smaller subtrees in Nextstrain. Similarly, Pathoplexus provides links to Nextstrain real-time analyses as a recognized resource for understanding the data housed within the repository, and other open-source software have used Nextstrain tools to aid in establishing new nomenclature systems for emerging pathogens (e.g., [26]). These examples illustrate how the development of open, flexible software can lead to interoperability across tools which further increases our ability to gain insights into pathogen dynamics.

## 5 Conclusion

Pathogen genomic surveillance has become a vital tool for enabling effective public health responses to emerging and endemic threats. Platforms such as Nextstrain have demonstrated the impact these tools can achieve when operating within a robust ecosystem of open data sharing among data generators and analysis groups. For such an ecosystem to function sustainably, resources must be shared in ways that maximize accessibility while ensuring appropriate credit to those who generate them. By fostering global collaboration among public health agencies and academic laboratories, genomic surveillance can provide critical situational awareness for future outbreaks, epidemics, and pandemics.

## Acknowledgements

We gratefully acknowledge the thousands of individuals who contributed to generating and sharing pathogen genomic sequences used in the real-time analyses. We also thank authors, originating laboratories, and submitting laboratories of these sequence data, as well as NCBI, Pathoplexus and GISAID for hosting databases and providing access to the data. We thank Allison Black, Vítor Borges, Sarah Cobey, Jason Caravas, Crystal Gigante, George Githinji, Peter van Heusden, Angie Hinrichs, Zamin Iqbal, Nicola Lewis, Stephanie Lunn, Placide Mbala, Laura McMullan, Marc Perry, Andrew Rambaut, Bryan Tegomoh, Sofonias Tessema, Pauline Trinh, Nídia Trovão, Mayra Trujillo, Erik Wolfson, Michael Zeller, and other public health workers and researchers for their feedback on these real-time pathogen analyses.

## Data availability

The sequence data and metadata used in the Nextstrain real-time analyses include all sequence data available for the 22 target pathogens in GenBank, SRA, and Pathoplexus, and all sequence data available for influenza and SARS-CoV-2 in GISAID. For data that are openly shared in these repositories, the sequence data and curated metadata are also made available through API access to data.nextstrain.org. A user interface to search these files is available at nextstrain.org/pathogens/files.

## Supplementary Information

**Supplementary Table 1.** Scaling of workflow run time and memory, and visualization performance across dataset sizes of SARS-CoV-2. This is running the standard “ncov” workflow at https://github.com/nextstrain/ncov to produce a single build target, namely ncov/open/global/all-time/. This uses 100k pre-filtered dataset to avoid overhead of subsampling from the ~9M publicly available genomes. All runs were on a single AWS c7a instance with 2 cores allocated to the workflow to equilibrate performance while providing enough memory for the large datasets to complete.

| Tips | Total wall | Tree | Refine | Peak RAM | JSON | Page load | Recolor |
| --- | --- | --- | --- | --- | --- | --- | --- |
| 500 | 4.4 min | 32 sec | 2.5 min | 0.6 GB | 1.0 MB | 144 ms | 63 ms |
| 1000 | 8.6 min | 1.6 min | 5.0 min | 0.9 GB | 1.9 MB | 190 ms | 108 ms |
| 2000 | 20 min | 6.9 min | 10 min | 2.1 GB | 3.5 MB | 267 ms | 216 ms |
| 5000 | 74 min | 39 min | 28 min | 7.3 GB | 7.6 MB | 487 ms | 420 ms |
| 10,000 | 3.6 hr | 2.3 hr | 1.0 hr | 17 GB | 15 MB | 842 ms | 603 ms |
| 20,000 | 11.2 hr | 8.6 hr | 2.2 hr | 45 GB | 28 MB | 1461 ms | 885 ms |

### Supplementary Text

#### Development of the Nextstrain ecosystem since 2018

The initial Nextstrain publication [1] described the platform shortly after its inception when it comprised the Augur analysis pipeline and the Auspice visualization application applied to a small number of centrally maintained pathogen builds. The modular Augur architecture that underlies the workflows described here was itself introduced at Augur version 3.0.0 in September 2018, and the ecosystem has since expanded substantially in both capability and scope. We provide a non-exhaustive summary of major developments below, organized by component.

### Augur

Augur was refactored from a largely monolithic prepare-and-process pipeline into a modular toolkit [7] of composable subcommands (e.g., `filter`, `align`, `tree`, `ancestral`) orchestrated by Snakemake, beginning at version 3.0.0 in September 2018. This composable design is the basis of the workflows described here. This toolkit saw continual feature additions with improvements to subsampling, merging data sources, VCF inputs, etc… A dedicated `augur curate` suite for standardizing and harmonizing metadata was introduced at version 18.2.0 in November 2022 and expanded substantially thereafter; this suite now underpins the metadata curation performed in the ingest workflow.

### Nextclade

Nextclade [9] did not exist at the time of the 2018 publication (version 1.0.0 was released in June 2021). It performs reference-based alignment, sequence quality control, clade and lineage assignment, mutation calling relative to a reference sequence and clade founders, and is available as both a web application and a command-line tool. Its development enabled the standardized quality-control and clade-calling step now incorporated into our ingest workflows.

### Auspice

Auspice has grown from simply a geographic map and phylogenetic tree viewer into a versatile analysis tool for phylodynamic data. New visualisations include a genomic diversity/entropy panel, clade frequency plots, second-tree and tanglegram views, and a measurements panel for displaying titer or deep mutational scanning data [46]. Auspice also gained Narratives, which pair explanatory prose with linked views into the underlying data that update automatically as the reader moves through the narrative [8]. These changes have necessitated a new v2 dataset schema shared with `augur export v2`.

### Nextstrain CLI and runtimes

The `nextstrain` command-line tool was developed to make installation and execution portable and reproducible across computing environments. It supports multiple runtimes, including Docker, Conda, ambient, and Singularity/Apptainer, as well as job submission to AWS Batch. This infrastructure supports the more computationally demanding workflows described here.

### nextstrain.org and data sharing

Capabilities for sharing data and results have expanded well beyond centrally hosted builds. Nextstrain Groups allow institutions and public health agencies to share datasets publicly or privately; Community sharing renders datasets directly from any GitHub repository; and a RESTful API at data.nextstrain.org provides programmatic access to curated sequences, metadata, and analysis outputs, including the per-pathogen data files now surfaced at nextstrain.org/pathogens/files. Finally, versioned dataset access is available for our core builds to enable retrospective viewing of analyses from any point in time.

### Standardized, automated pathogen workflows

The standardized per-pathogen repository structure that is the primary subject of this paper was developed since our prior publication. Key features of this structure include separate ingest, Nextclade, and phylogenetic Snakemake workflows, shared curation tooling, and GitHub Actions automation producing continually updated results. The separation of individual pathogen builds into dedicated repositories began in September 2018, and the automated, continually updated workflow architecture described here has been built out in the years since.

